# Semantic relatedness emerges in deep convolutional neural networks designed for object recognition

**DOI:** 10.1101/2020.07.04.188169

**Authors:** Taicheng Huang, Zonglei Zhen, Jia Liu

## Abstract

Human not only can effortlessly recognize objects, but also characterize object categories into semantic concepts and construct nested hierarchical structures. Similarly, deep convolutional neural networks (DCNNs) can learn to recognize objects as perfectly as human; yet it is unclear whether they can learn semantic relatedness among objects that is not provided in the learning dataset. This is important because it may shed light on how human acquire semantic knowledge on objects without top-down conceptual guidance. To do this, we explored the relation among object categories, indexed by representational similarity, in two typical DCNNs (AlexNet and VGG11). We found that representations of object categories were organized in a hierarchical fashion, suggesting that the relatedness among objects emerged automatically when learning to recognize them. Critically, the emerged relatedness of objects in the DCNNs was highly similar to the WordNet in human, implying that top-down conceptual guidance may not be a prerequisite for human learning the relatedness among objects. Finally, the developmental trajectory of the relatedness among objects during training revealed that the hierarchical structure was constructed in a coarse-to-fine fashion, and evolved into maturity before the establishment of object recognition ability. Taken together, our study provides the first empirical evidence that semantic relatedness of objects emerged as a by-product of object recognition, implying that human may acquire semantic knowledge on objects without explicit top-down conceptual guidance.

**Significance Statement:** The origin of semantic concepts is in a long-standing debate, where top-down conceptual guidance is thought necessary to form the hierarchy structure of objects. Here we challenged this hypothesis by examining whether deep convolutional neural networks (DCNNs) for object recognition can emerge the semantic relatedness of objects with no relation information in training object datasets. We found that in the DCNNs representations of objects were organized in a hierarchical fashion, which was highly similar to WordNet in human. This finding suggests that top-down conceptual guidance may not be a prerequisite for human learning the relatedness among objects; rather, semantic relatedness of objects may emerge as a by-product of object recognition.

## Introduction

Objects in this world is complicated. Variations of object (e.g. orientation, size, shape and color) create challenges for human to flexibly recognize and categorize them (Logothetis et al., 1996). To survive in such difficult and diverse environment, human learn to characterize objects into a rich and nested hierarchical structure, which finally evolves into semantic concepts (Tanaka, 1996; Yamins et al., 2014). However, how the hierarchically-structured semantic concepts are formed is still under hotly debated.

Two hypotheses have been proposed. One hypothesis (Leshinskaya & Caramazza 2016; Mahon & Caramazza, 2009) suggests that semantic concepts is only formed and accessed through abstract symbols that are independent of perceptual experiences. Supporting evidence comes from studies on congenitally blind people, whose core semantic retrieval system in the frontal-temporal cortex can still be activated for retrieving visually-experienced semantic information (Noppeney et al., 2003; Noppeney et al., 2007). In addition, functional brain imaging studies find that supramodal regions in ventral temporal occipital cortex (e.g. superior occipital, inferior and superior parietal areas) are also involved in processing objects in blind individuals (Lambert et al., 2004; Riccardi et al., 2014). Therefore, perceptual experiences seemed not necessary for the emergence of semantic concepts.

An alternative hypothesis argues that the development of semantic concepts derives from the perception of natural environment (Barsalou, 2008; Roy, 2005; Sloutsky, 2003). For example, in a word/no word match-to-sample task, Imai et al. (1994) decouple taxonomic and perceptual similarity of words, and find that younger children rely on the visual property of objects, rather than taxonomic concepts, in response to novel words. A more direct evidence comes from a study on 10-month-old infants who learn new words by the perceptual salience of an object rather than social cues provided by the caregivers (Pruden et al., 2006). That is, perceptual features are needed to form semantic concepts.

One inevitable limitation of these studies is that perceptual experiences and conceptual guidance are tightly intermingled during the development; therefore, it is impossible to examine one factor with the other controlled. In contrast, the advance of deep convolutional neural networks (DCNNs) provides a perfect model to examine how semantic relatedness is formed. On one hand, DCNNs have abundant visual experiences on objects, as with the presence of millions of natural images, the DCNNs learn to extract critical visual features to classify objects into categories as perfectly as human. On the other hand, during the training, the relation among object categories is not provided in the training task or in the supervised feedback. Therefore, conceptual guidance is completely absent in the DCNNs. With such characteristics of the DCNNs, here we asked whether semantic relatedness among object categories was able to emerge with no top-down conceptual guidance.

To address this question, we used two typical DCNNs, AlexNet and VGG11, which are designed for classifying objects into 1,000 categories in ImageNet. Specifically, we first measured whether the representations of some object categories were more similar than their relation to others, which formed a hierarchical structure of object categories as a whole. We reasoned that if a stable and well-organized hierarchical structure was observed, the hypothesis of the necessity of conceptual guidance in forming the semantic relatedness was challenged.

### Materials and Methods The ImageNet dataset

We used the ILSVRC2012 dataset as the image stimulus (http://image-net.org/challenges/ILSVRC/2012/). Both training and validation datasets were used in this study. The ILSVRC2012 training dataset contains about 1.2 million images with labels from 1,000 categories. The validation dataset contains 50,000 unduplicated images that belong to the same 1,000 categories as the training dataset.

The sets of synonyms (synsets) of the 1,000 categories were followed the hierarchical structure of WordNet (Miller, 1995). Among the 1,000 synsets, 3 synsets belong to abstract entity (bubble, street sign, and traffic light), 39 synsets to matter, 9 synsets to geological formation, 517 synsets to the artifact, 407 synsets to the living thing and 16 synsets to fruit (Figure 1A). All categories have an ontology root (e.g. entity) and most of them are subsets of the physical entity (997 categories). Additionally, all of the 1,000 synsets have no parent-child relationship with each other (Russakovsky et al., 2015).

**Figure 1.**
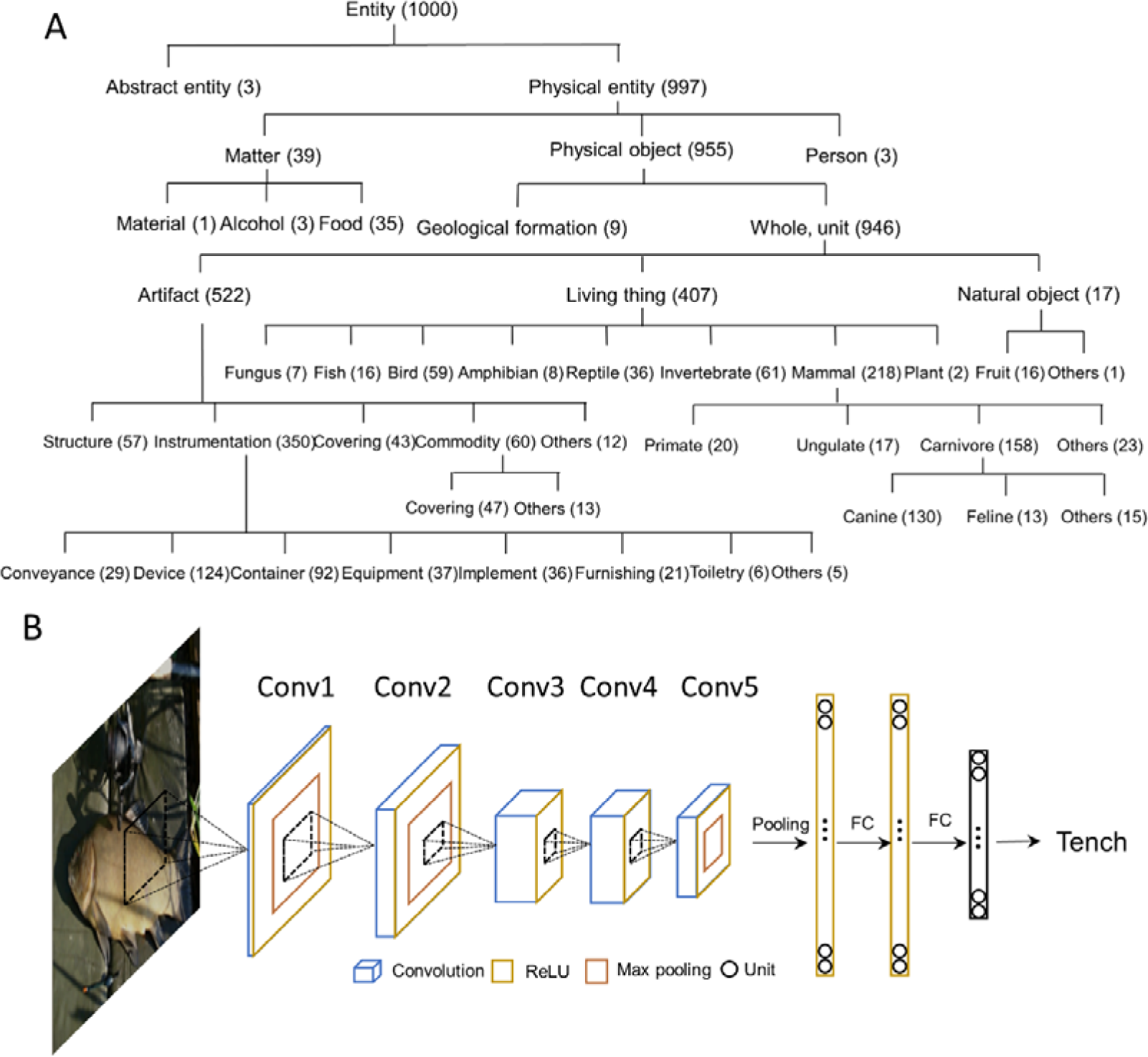
(A) The WordNet hierarchical structure of the 1,000 categories. All categories were derived from an ontology root (e.g. entity), and most of them are subsets of physical entity. The 1,000 categories cover a wide range of physical objects, make it suitable to represent the conceptual relationship. Numeral after each word are the number of categories belonging to this superordinate category. (B) **The architecture of AlexNet**. The AlexNet includes 8 layers of computational units stacked into a hierarchical architecture: the first 5 are convolutional layers, and the last 3 layers were fully connected for category classification.

### Deep convolutional neural networks (DCNNs)

Two fully-pretrained DCNNs, AlexNet (Krizhevsky et al., 2012) and VGG11 (Simonyan & Zisserman, 2014), were used in this study, which were pretrained on ImageNet for classification of 1,000 object categories. The models were downloaded from PyTorch model Zoo (https://pytorch.org/docs/stable/torchvision/models.html). AlexNet’s architecture includes 8 layers of computational units stacked into a hierarchical architecture: the first 5 are convolutional layers, and the last 3 layers are fully-connected for category classification (Figure 1B). ReLU nonlinearity is applied after all layers except for the last fully-connected layer. The first two and the last convolutional layers are followed by the max-pooling layers while the third and fourth convolutional layers are connected directly. Layer 1 through 5 consists of 64, 192, 384, 256 and 256 kernels. Layer 6 to 8 consist of 4,096, 4,096 and 1,000 units which were fully connected. The top-1 and top-5 accuracies of AlexNet are 56.5% and 79.1%.

VGG11’s architecture includes 11 layers that stacked into a hierarchical structure, the first 8 are convolutional layers, and the last 3 are fully-connected layers (Supplementary Figure 1). ReLU nonlinearity was applied after all layers except for the last fully-connected layer. Max-pooling layers then followed the 1, 2, 4, 6, 8 convolutional layers while others are connected directly. Layer 1 through 8 consisted of 64, 128, 256, 256, 512, 512, 512 and 512 kernels. For classification, the number of units in layer 8 to 11 were 4,096, 4,096 and 1,000, respectively. The top-1 and top-5 accuracies of VGG11 are 69.0% and 88.6%.

### The semantic similarity of category in WordNet

The semantic similarity of the 1,000 categories was evaluated by the WordNet 3.0 (Miller, 1995), which is one of the most popularly-used and largest lexical databases of English. In WordNet, categories are organized into synsets, each representing a lexicalized concept. The lexical hierarchy is connected by several superordinate terms in semantic relations, providing a hierarchical tree-like structure for each term. All of 1,000 categories were separately extracted for generating a hierarchical tree-like structure by referring to their WordNet IDs.

We measured the semantic similarity between each pair of the 1,000 categories using Wu and Palmer’s similarity (Wu & Palmer, 1994), which computed the similarities between concepts in an ontology restricted to taxonomic links. This measure is given by:

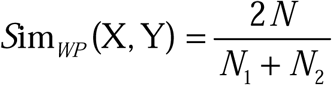

Where N_1_ and N_2_ are the depth between the concepts X, Y and the ontology root (i.e. ‘entity’ in WordNet) and N is the depth between the least common subsume (i.e. most specific ancestor node) and the ontology root.

### Representation similarity of categories in DCNNs

Responses to each image were extracted from all of the convolutional layers and the last fully-connected layer using the DNNBrain toolbox (https://github.com/BNUCNL/dnnbrain). No ReLU was performed for the responses. Responses of stimulus from the same category were averaged to make a response pattern for this category. The category similarity of a layer was measured as correlation of response patterns between each of two categories. In addition, correspondence between the category representational similarity from the DCNNs and the WordNet semantic similarity was calculated to measure the extent to which the relatedness of objects in the DCNNs was similar to that in human.

### The development of the relatedness in DCNNs

To investigate how the hierarchical structure of objects emerged in the DCNNs, we separately retrained the AlexNet and VGG11 from scratch with about 1.2 million images that belong to the 1,000 categories from the ImageNet training dataset (Deng et al., 2009) using the PyTorch toolbox (Paszke et al., 2019). The network was trained for 50 epochs, with the initial learning rate as 0.01 and a step multiple of 0.1 every 15 epochs. The parameters of each model were optimized using stochastic gradient descent with the momentum and weight decay was fixed at 0.9 and 0.0005, respectively. Each input image was transformed by random crop, horizontal flip and normalization to improve the training effect of the network. During the training progression, object classification accuracy was evaluated in predicting the category of 50,000 images that from the ILSVRC2012 validation dataset in each epoch. In the end, the top-1 and top-5 accuracies for the AlexNet were 51.0% and 74.5%, and for the VGG11 were 63.2% and 85.0%, which were comparable to the pretrained models from the PyTorch model Zoo.

During the training progression, we input images from the ILSVRC2012 validation dataset by simply feedforwarding in each epoch to get the activation responses, and then averaged responses within each category and computed the similarity between each pair of categories for the category similarity. Correspondence between the category similarity from the DCNNs and the WordNet semantic similarity in each training stage was measured to evaluate how similar the relatedness of objects was between the DCNN and human.

To reveal at which semantic level the category similarity from the DCNN showed better correspondence to the WordNet semantic similarity, the category similarity from the DCNN was measured at coarse level and fine-grained level, respectively. In particular, we first manually selected 19 superordinate concepts (i.e. food, fungus, fish, bird, amphibian, reptile, mammal, invertebrate, conveyance, device, container, equipment, implement, furnishing, toiletry, covering, commodity, structure and geological formation) that covered most of the 1,000 categories by referring to the WordNet hierarchical relationship, then grouped categories into these superordinate concepts. The coarse-grained correspondence was measured as the correlation between the DCNNs category similarity and the WordNet semantic similarity in 19 superordinate concepts. In turn, the similarity among superordinate concepts were calculated by averaging the category representation similarities from each pair of superordinate concepts. The fine-grained correspondence was measured as the averaged correspondence between the DCNNs category similarity and the WordNet semantic similarity within each superordinate concept.

### Data availability

All data and analysis scripts were available for download at https://github.com/helloTC/SemanticRelation.

## Results

We first evaluated whether there was a hierarchical structure among object categories in the AlexNet, which was trained to classify object categories from the ImageNet containing no relation information among objects. For this, responses from the last fully-connected layer of the AlexNet (i.e., FC3) were averaged across images of each category as the response pattern for this category, and the similarity between two categories was calculated as the correlation between their response patterns. A great variance in similarity was observed, with the highest similarity between object toy poodle and object miniature poodle (r = 0.99), the lowest between object snail and object fur coat (r = -0.62), and the mean similarity of r = 0.21. The variance in similarity observed was significantly larger than variance from a randomized structure (permutation analysis, p < .001), suggesting that objects were structurally organized (Figure 2A, left). A close inspection of Figure 2A revealed two large clusters, one is animate things and the other artefacts. Within each cluster, there are sub-clusters, as within-cluster variance was smaller than that of neighboring sub-clusters. The nested structure in similarity suggests that a hierarchical relation among objects emerged in the AlexNet without conceptual guidance.

**Figure 2.**
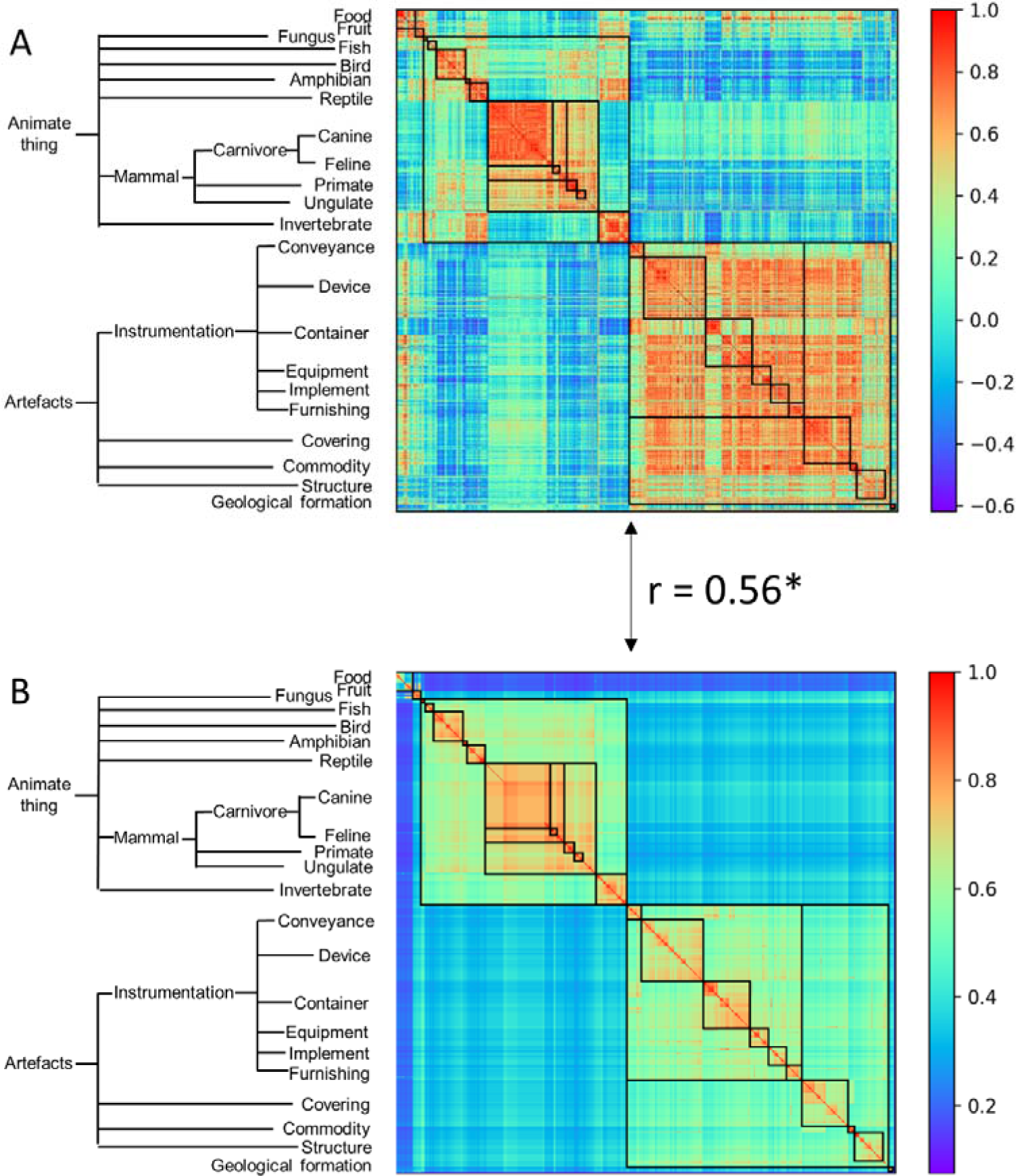
The category representational similarity of the AlexNet (A) and the semantic similarity of WordNet hierarchy (B). Categories were ordered according to the WordNet semantic hierarchy. A simplified hierarchical structure was shown as indicator of superordinate categories in WordNet semantic similarity. For the ease of comparison between AlexNet’s category similarity and WordNet semantic similarity, categories belong to the same superordinate category were marked with a black box. The AlexNet category similarity showed good correspondence to the WordNet semantic similarity. Asterisk denotes p < 0.001.

A similar nested structure of objects was also observed in the VGG11, which is designed for the same task but with different network architecture (e.g., layer number and kernel size) (Supplemental Figure 2A). Importantly, the hierarchical relation of the object categories emerged from the VGG11 was almost identical to that from the AlexNet (r=0.96), implying that the emerged hierarchical relation among object categories was invariant to implementations, but rather result from inherent properties of stimulus and the task that DCNNs received and performed. Because human brains used images from the same physical world to perform the same task, an intuitive thought is that the hierarchical relation observed in the DCNNs may be similar to semantic relatedness of objects in human.

To test this conjecture, the names of the object categories were put into WordNet derived from human, and their semantic similarity was calculated with the Wu and Palmer’s similarity approach (Figure 2B). We found that there was a significant correlation between semantic similarity of WordNet in human and the hierarchical relation among objects in the AlexNet (r = 0.56, p<.001), and correlation also reached significance for both the animate thing (r = 0.70, p<.001) and artefact (r = 0.41, p<.001). In addition, the correspondence increased as a function of layers (Figure 3), with lower correlations observed in first two layers (layer 1: r = 0.21, layer 2: r = 0.15) and higher correlations in the third (r = 0.41), forth (r = 0.51) and fifth (r = 0.53) layers. A close inspection on the increases of hierarchy among layers revealed that coarse structure (e.g., animates versus artefacts) was first emerged in lower layers, and a fine-grained structure was prominent only in higher layers.

**Figure 3.**
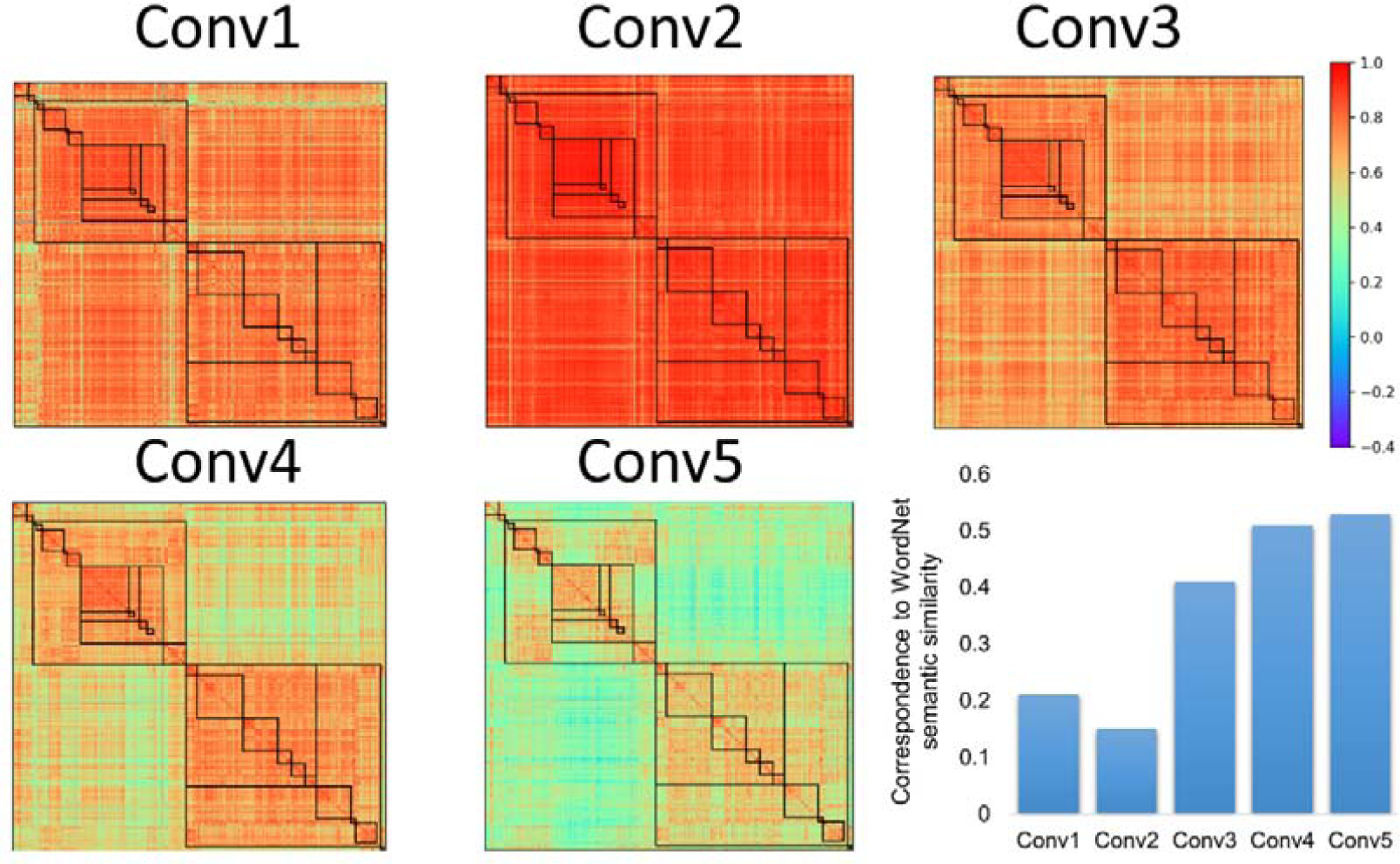
The category representational similarity in different convolutional layers of the AlexNet. Hierarchical relations of objects in the AlexNet gradually emerged as a function of convolutional layers, so was the correspondence between the representational similarity in the AlexNet and WordNet semantic similarity in human. Coarse structure first emerged in lower layers, while fine-grained structure was prominent only in higher layers.

How did the hierarchical relatedness of object categories emerge from unstructured image dataset in the DCNNs? To address this question, we explored the developmental trajectory of the relatedness when the AlexNet was trained to recognize objects. A new AlexNet was trained from scratch for 50 epochs to achieve comparable top-1 and top-5 accuracies as the pretrained AlexNet. Two findings were observed. First, the correspondence in the hierarchical relatedness of object categories between the AlexNet and the WordNet was established within the first epoch (r = 0.60, Figure 4A), whereas the performance for object recognition (top-1 accuracy: 8.9%) was far below that of the fully trained one (top 1 accuracy: 51%). Instead, at least 40 training epochs were needed to attain the matched performance to the fully trained model. The asynchronous development illuminated that the relatedness of object categories in the AlexNet was formed before it was capable of performing the task. Second, within the development of the hierarchical relatedness, there was a progression from a coarse structure to a fine-grained structure. That is, the coarse structure based on the 19 concepts (e.g., bird and device) merged from 1,000 object categories reached a plateau within the first epoch (Figure 4B), with a correlation of 0.65 to the WordNet. In contrast, the fine-grained structure within the 19 concepts (e.g., crane and flamingo in bird) did not approach a plateau until 40 epochs’ training, with an averaged correlation of 0.38 to WordNet in human. Similar developmental trajectory was also observed in the VGG11 (Supplemental Figure 3). Therefore, the hierarchical relatedness of object categories was formed in a coarse-to-fine fashion, with the coarse structure formed before the fine-grained structure.

**Figure 4.**
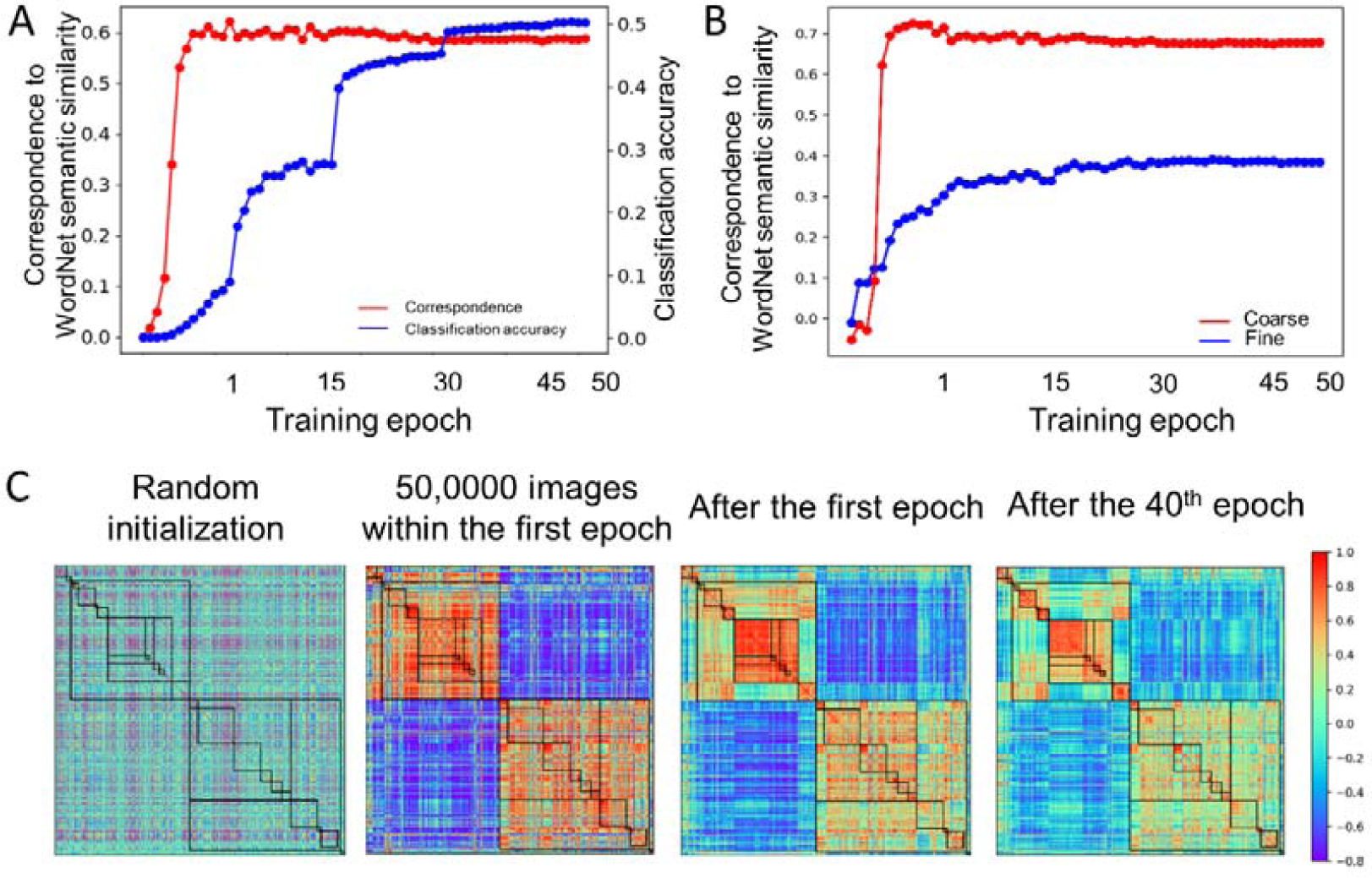
Developmental trajectory of the relatedness. (A) The developmental trajectory of the correspondence in the hierarchical relatedness of object categories between the AlexNet and WordNet (red line). The classification accuracy of the AlexNet was shown in blue. The hierarchical structure evolved into maturity far prior to the establishment of object recognition ability. (B) A coarse to fine shift during training progression. The coarse structure based on the 19 superordinate categories reached a plateau within the first epoch (red line), while the fine-grained structure reached a plateau after 40 epochs’ training (blue line). (C) The category similarities of the AlexNet in different training stages for comparison. From left to right, category similarities of the AlexNet without training, AlexNet trained with 50,0000 images within the first epoch, AlexNet trained after the first epoch and AlexNet trained after the 40^th^ epoch. Color bar indicates correlation coefficients.

## Discussion

In this study, we used DCNNs as a model for human cognition to examine whether the semantic relatedness of object categories can automatically emerge without top-down conceptual guidance. First, we found that an almost identical hierarchical structure of object categorizes emerged in both AlexNet and VGG11, which was highly similar to WordNet derived in human. This result suggests that the relation among object categories can be automatically formed without conceptual guidance and independent of implementation hardware. Further investigation on the developmental trajectory of the relatedness revealed that it was constructed prior to the ability of object recognition in a coarse-to-fine fashion. In sum, our study provided the first empirical evidence that the semantic relation of objects can be formed from perceptual experiences alone.

Unlike studies on human where perceptual experiences are always intermingled with conceptual guidance, DCNNs provide a perfect model to demonstrate that the perceptual experience alone was sufficient to construct the relatedness among objects. This finding is in line with developmental studies in which children prefer naming objects by referring their perceptual features, suggesting that the perceptual property of objects play an important role in accessing lexical knowledge (Gershkoff-Stowe and Smith, 2004; Samuelson and Smith, 2005). Further, the effect of perceptual experience is likely independent of implementation, because the DCNNs and human brain, which differ significantly in hardware, shows highly similar hierarchical structure of objects. The similarity in the semantic structure may result from the similarity in architecture that DCNNs are designed with an architecture similar to human visual cortex. Accordingly, similar anatomy may lead to similar structure of the relatedness among objects. Indeed, the top level of the hierarchy was the animate things versus artefacts, mirroring the axis of the mid-fusiform sulcus that separates the coding of animate objects and artefacts in the brain (Kalanit and Kevin, 2014).

Another and more plausible possibility may be the way by which objects are coded in a representational space. In DCNNs, an object is firstly decomposed into multiple features, and mapped to a representational space. Then, the object is reconstructed from the feature repertoire of the representational space based on the demand of tasks (Xiang et al., 2019; Yang et al., 2019). Further, features of the representational space are distributedly represented by different units (Yang et al., 2020); therefore, if two objects are perceptually similar because of shared features, they are likely represented by the same set of units. In this way, the relation between two objects is then derived from the connections among units. In fact, this intuition is consistent with the hypothesis of parallel distributed processing (McClland and Rogers, 2003; see Saxes et al., 2019 for mathematical theory), where knowledge arises from the interactions of units through connections. Accordingly, the knowledge stored in the strengths of the connections finally becomes the building blocks of the hierarchical structure of object categories.

Importantly, such hierarchical structure emerged in a coarse-to-fine fashion. That is, at the initial stage of learning, DCNNs may encode global features to identify relations among objects when only a small number of exemplars are available. For example, dogs and cats are the same, but they are not trees based on general appearance. When more exemplars are learned, features in the repertoire are greatly enriched, and thus are capable of providing fine-grained representations for objects to establish the hierarchical structure of relation among objects. This coarse-to-fine representation is also observed in infants, as infants are able to distinguish animals and vehicles at 7 months old, but fail to differentiate dogs from cats until 11 months old (Mandler and McDonough, 1993; Mandler and McDonough, 1998; also see Pauen, 2002).

Interestingly, we also found that the hierarchical structure evolved into maturity before the establishment of object recognition ability. This is not surprising because the enriched and structured feature repertoire is necessary for DCNNs to successfully recognize novel objects never seen before. For example, in a recent study where DCNN’s experience on faces is selectively deprived, and the DCNN is still capable of successfully accomplishing a variety of faces tasks behaviorally and evolving face-specific modules internally (Xu et al., 2020). Therefore, a mature representational space of objects will greatly benefit DCNNs’ performance. This mechanism has already widely used in computer science, as transfer learning, for example, utilize it to harness a pretrained network to work in another domain with a small number of exemplars but still with a high accuracy (Torrey and Shavlik, 2010).

In sum, our study demonstrated that perceptual similarity among object categories alone was sufficient for DCNNs to construct hierarchical structure among objects. However, this finding does not necessarily rule out the role of conceptual guidance in forming the semantic relatedness, which is clearly illustrated by a moderate correlation between the DCNNs and human in the hierarchical structure among objects. In addition, the DCNNs used in this study are purely feedforward, and may not be suitable for studies on conceptual guidance. Therefore, other deep neural networks with feedback connections, such as Feedback-CNN or predictive coding network (Cao et al., 2015; Lotter et al., 2016), or networks directly trained with lexical and semantic constraints (Bayer and Riccardi, 2015), shall be used to understand how conceptual guidance modulates the semantic relatedness of objects without the influence of perceptual experiences.

## Acknowledgements

This study was funded by the National Natural Science Foundation of China (Grant No. 31861143039, 31771251), the National Key R&D Program of China (Grant No. 2019YFA0709503), and the National Basic Research Program of China (Grant No. 2018YFC0810602).

