## Supplementary Materials for "Semantic relatedness emerges in deep convolutional neural networks designed for object recognition"

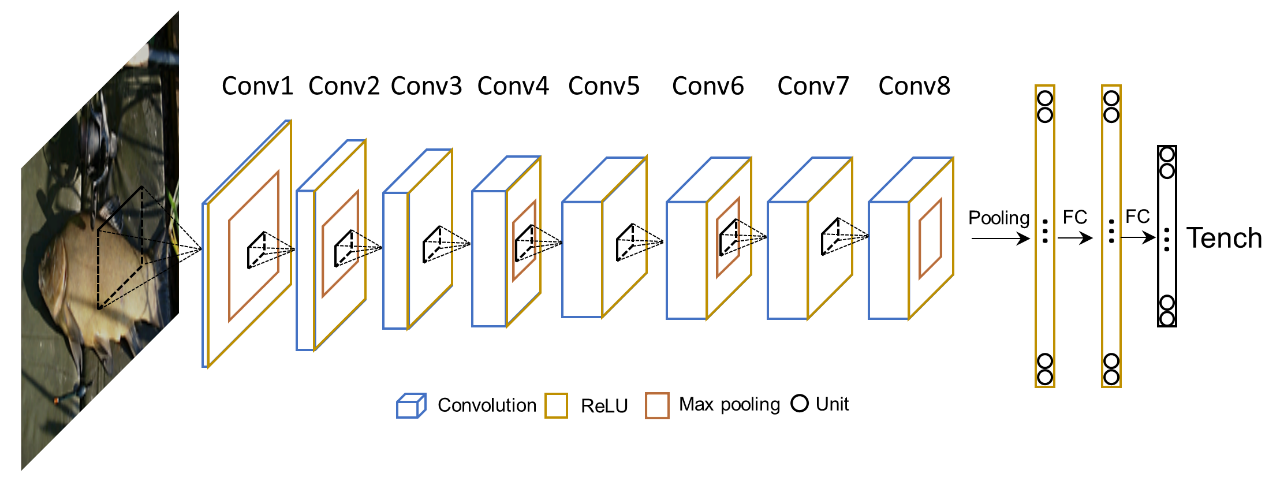

Supplementary Figure 1 **The architecture of VGG11**. VGG11 included 11 layers of computational units stacked into a hierarchical architecture: the first 8 were convolutional layers, and the last 3 layers were fully connected for image recognition classification.

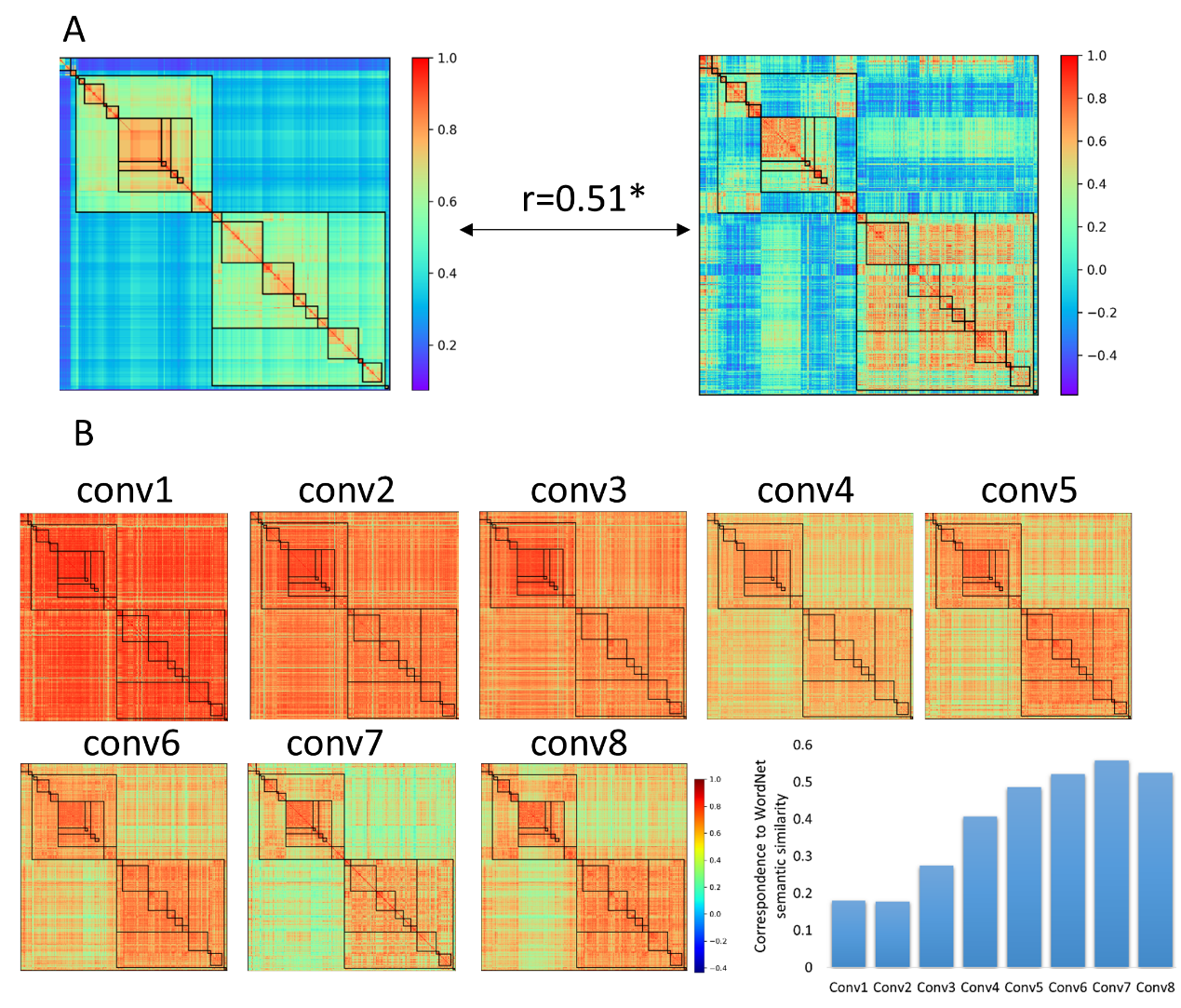

Supplementary Figure 2 (A) **Hierarchical relation emerged in the VGG11.** The category representational similarity of the VGG11 (left) and the semantic similarity of the WordNet hierarchy (right). The VGG11 category similarity showed good correspondence to the WordNet semantic similarity as that from AlexNet. Asterisk indicates p < 0.001. (B) **The category representational similarity of the VGG11 in different convolutional layers.** The correspondence increased as a function of convolutional layers.

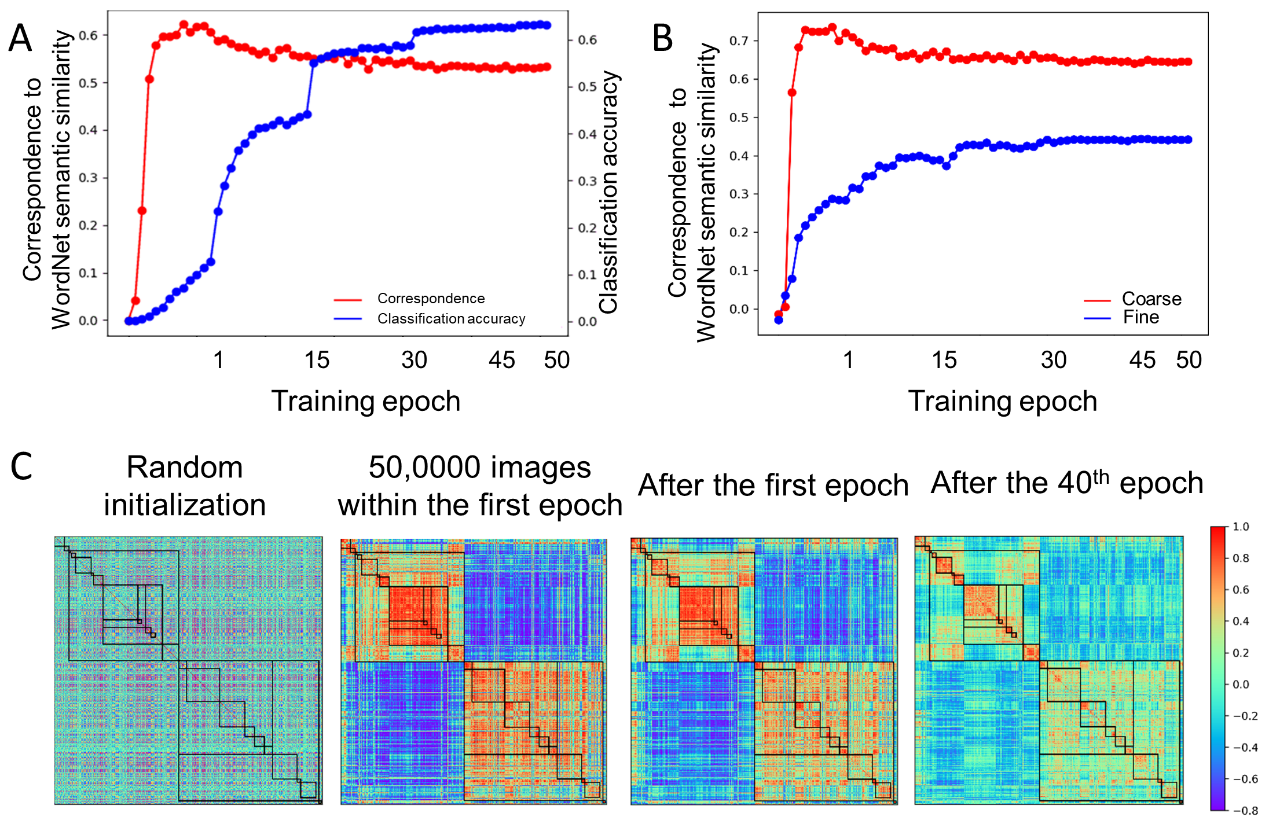

Supplementary Figure 3 **Developmental trajectory of the semantic relatedness when the VGG11 was trained to recognize objects.** A) The correspondence in the hierarchical relatedness of object categories between the VGG11 and the WordNet was established within the first epoch (red line), while classification accuracy (blue line) was far below than that of the fully trained model. B) A coarse to fine shift was observed during the training progression. The coarse structure based on the 19 superordinate categories reached a plateau within the first epoch (red line) while fine-grained structure reached a plateau after 40 epochs’ training (blue line). C) The category similarities of the VGG11 in different training stages for comparison. From left to right, category similarities of the VGG11 without training, VGG11 trained with 50,0000 images within the first epoch, VGG11 trained after the first epoch and VGG11 trained after the 40th epoch. Color bar indicates correlation coefficients.
